# Mechanical signatures of nucleic acid knot topology

**DOI:** 10.64898/2026.03.16.712236

**Authors:** David t.R. Bakker, Micah Yang, Isaac T.S. Li

## Abstract

Molecular knots represent a fundamental form of polymer topology, yet their mechanical behaviour in nucleic acids remains largely unexplored. Here, we engineer a single-stranded DNA sequence that can fold into either a knot or a pseudoknot while maintaining identical base-pairing interactions. Using single-molecule force spectroscopy with optical tweezers, we show that molecular topology alone produces distinct mechanical behaviour. Knotted ssDNA exhibits three characteristic signatures relative to the pseudoknot: higher unfolding forces, shorter unfolding extensions, and faster refolding kinetics. These features arise from the topological constraint imposed by strand threading and together provide a mechanical fingerprint that distinguishes knotted from unknotted nucleic acid structures. By analyzing the denatured state under tension, we further show that the knot tightens as force increases, entering a tight-knot regime in which the molecule can be described as a compact knot core in series with a stretched single strand. The retained contour length reveals that the tight trefoil knot contains approximately ten nucleotides at forces approaching 40 pN. These results establish mechanical signatures as a means of identifying nucleic acid topology and provide quantitative insight into the nanomechanics of molecular knots.

## Introduction

Biological macromolecules can adopt topologically complex structures that influence their stability and function. Among these, molecular knots represent one of the most striking topologies accessible to polymer chains. Knots arise spontaneously in long polymers such as double-stranded DNA (dsDNA) through thermal fluctuations^1^, as well as from enzymatic processes in viruses^2^, bacteria^3^, and eukaryotes^4,5^. In dsDNA, these knots are typically loose and diffuse freely along the DNA contour without a fixed position. Excessive dsDNA knotting can disrupt genome organization and transcriptional processes, so cells employ topoisomerases to continuously remove these topological entanglements^6^.

In contrast to the transient knots observed in dsDNA, knotted topologies can stabilize folded biopolymers. Approximately 1% of known proteins contain topological knots in their native structures^7^. In these proteins, intramolecular interactions between amino acid side chains fix the knot at a specific location along the polypeptide chain. The defining step of knotted protein folding is the threading of a polypeptide terminus through a loop in a partially folded structure^8,9^, a kinetically and entropically unfavourable event that determines the topology of the final fold. Once formed, the knotted topology enhances protein thermodynamic stability and mechanical resistance to unfolding and degradation^10–15^.

Single-stranded nucleic acids (ssNA) can also adopt knot topologies through intramolecular base pairing and strand threading. Synthetic DNA and RNA knots have been engineered using rational sequence design and DNA nanotechnology approaches^16–21^. In these systems, the topology of the molecule was typically fixed by ligating the strand ends to form closed loops, allowing different knot topologies to be separated using gel electrophoresis. More recently, single-stranded DNA (ssDNA) topology has been examined without circularizing the molecule by ligating the strand ends to long dsDNA handles that kinetically lock the topology while enabling single-molecule force spectroscopy^22^. Despite these advances in synthetic design, naturally occurring nucleic acid knots remain extremely rare, with only a single RNA knot identified to date through structural biology techniques^23^.

Nucleic acid structures that appear topologically complex often instead fold into pseudoknots, a closely related but fundamentally different topology in which a true topological threading does not occur^24,25^. When mechanical force unfolds a pseudoknot from its ends, disruption of base pairing allows the structure to fully unravel. In contrast, a true knot retains its topological crossings even after all secondary structure is disrupted and therefore tightens under tension rather than disappearing, suggesting that mechanical measurements could distinguish the two topologies. This distinction raises a fundamental question: what are the nanomechanical signatures of a nucleic acid knot? Understanding how knots alter the mechanical response of nucleic acids is important for determining how molecular motors or nucleic acid processing enzymes interact with such topologies. However, direct experimental characterization of nucleic acid knots under force has remained limited.

Here, we engineer a single DNA sequence that can be folded into either a true knot or a pseudoknot while maintaining identical base-pairing interactions. Using single-molecule force spectroscopy with optical tweezers, we show that molecular topology alone produces distinct nanomechanical behaviour. Knotted ssDNA exhibits three characteristic mechanical signatures: increased unfolding force, reduced unfolding extension, and accelerated refolding kinetics. These signatures arise purely from the topological constraint imposed by the knot. Finally, by analyzing the force-dependent extension of the denatured state, we directly measure the tightening of the knot under tension and estimate the size of a trefoil knot in single-stranded DNA. Together, these results demonstrate that single-molecule force spectroscopy can identify nucleic acid topology and provide quantitative insight into the mechanics of molecular knots.

## Main

### Engineering sequence-identical ssDNA knot and pseudoknot

We designed a ssDNA sequence that can fold into either a knot or a pseudoknot while maintaining identical base-pairing interactions. The sequence contains two duplex-forming segments separated by poly(T) spacers that act as flexible loops (Fig. 1a). Each duplex-forming segment is 12 base pairs (bp) long, corresponding to just over one helical turn of B-form DNA. This geometry allows one strand of the DNA duplex to thread through its hairpin loop, forming a topological knot. The two duplex segments were designed with different base compositions: one CG-rich and one AT-rich. This difference creates distinct thermodynamic and mechanical stabilities, encouraging the two duplexes to form sequentially during folding. The poly(T) spacers were chosen to be sufficiently long to sterically permit full hybridization of the duplex segments while allowing strand threading during knot formation. Because both constructs share the same primary sequence and base-pairing interactions, this design isolates topology as the only structural variable.

**Figure 1:**
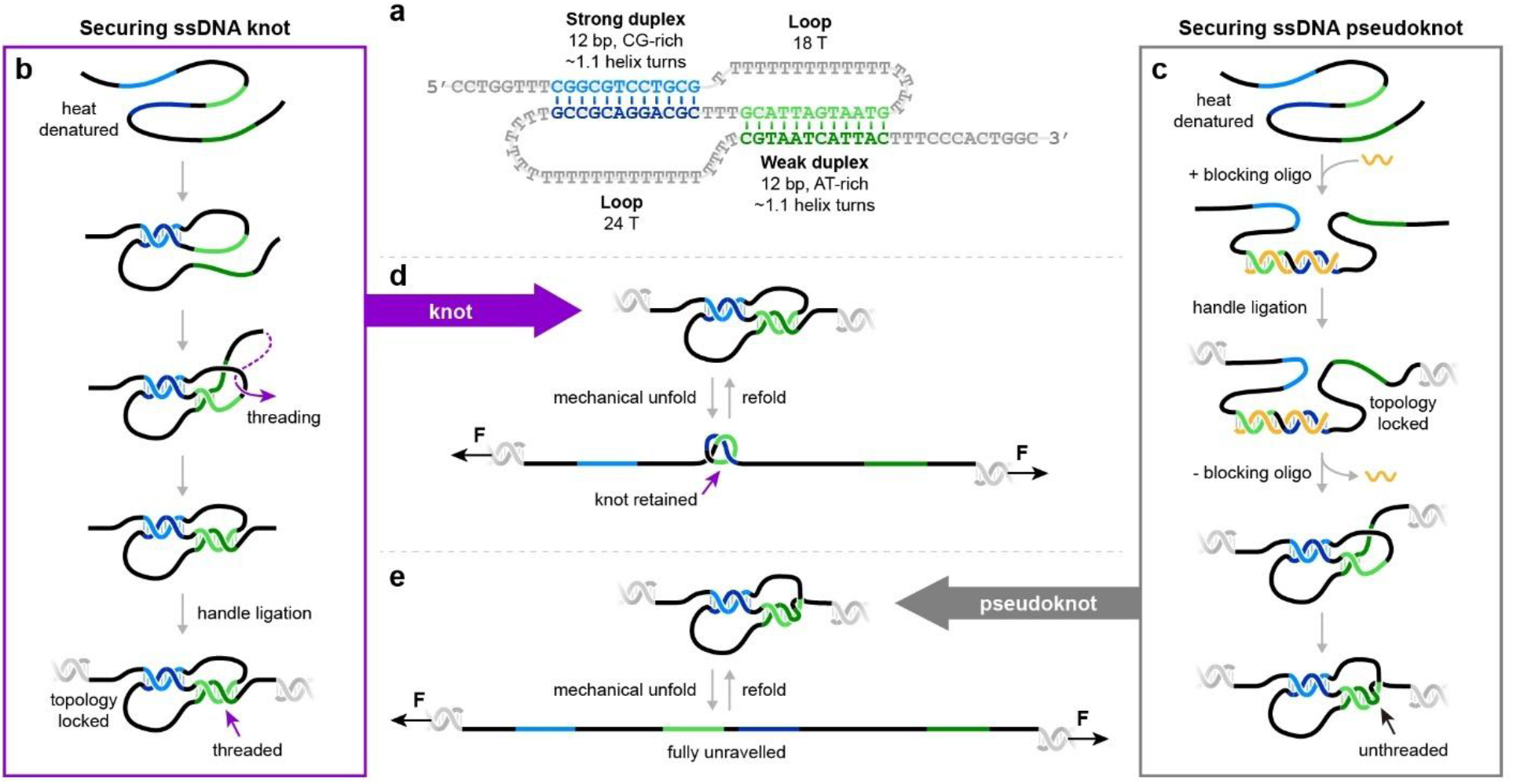
Construction of sequence-identical ssDNA knot and pseudoknot. **(a)** Design of the ssDNA construct. The sequence consists of two duplex-forming segments: a strong CG-rich duplex (blue, 12 bp) and a weak AT-rich duplex (green, 12 bp), separated by poly(T) loops. The 5’ and 3’ termini contain sticky ends for ligation to dsDNA handles. Note that the schematic represents base-pairing interactions but not topology. **(b)** Formation of the ssDNA knot. Heat-denatured ssDNA is slowly cooled to allow sequential duplex formation, where the strong duplex hybridizes first. Weak duplex formation subsequently drives free-end threading, forming a knot. Ligation of long dsDNA handles at both ends kinetically traps the threaded topology, securing the knot. **(c)** Formation of the ssDNA pseudoknot. After heat denaturation, a blocking oligonucleotide binds the duplex-forming regions to prevent self-threading. dsDNA handles are ligated while the strand is held in an unthreaded topology. Removal of the blocking oligo allows the strand to fold into a pseudoknot since the handles prevent knot formation. **(d, e)** Mechanical unfolding of the knot (d) and pseudoknot (e). Under tension, the pseudoknot fully unravels, while the knot retains its topological crossings, resulting in a shorter unfolded molecule.

To favour formation of a knotted topology, the ssDNA was thermally denatured and then slowly cooled (Fig. 1b). During cooling, the stronger CG-rich duplex forms first, creating a hairpin-like structure with a loop large enough to accommodate strand threading. Formation of the weaker AT-rich duplex then requires the free end of the strand to pass through this loop before hybridization can complete. The free energy gained from duplex formation drives this entropically and kinetically unfavourable threading event, producing a knotted configuration in which all 24 bp are hybridized. Long dsDNA handles, several kilobase pairs (kbp) in length and significantly more rigid than ssDNA, were subsequently ligated to both ends of the molecule. These handles kinetically trap the local topology, preventing the knot from diffusing out of the ssDNA region. In this work, the term “knot” refers to such open-ended knots, in which the topological entanglement is locally confined rather than formed by circularization of the strand.

The same ssDNA sequence can also be directed to form a pseudoknot by preventing strand threading prior to handle ligation (Fig. 1c). During thermal denaturation and slow cooling, a blocking oligonucleotide was added in excess that binds across both duplex-forming segments with a higher melting temperature than either duplex alone. This blocking strand suppresses spontaneous self-threading during folding. While the ssDNA remains bound to the blocking strand, the dsDNA handles are ligated to both ends of the molecule, thereby fixing the topology in an unknotted state. After ligation, the blocking strand is removed, allowing the ssDNA to fold. Under these constraints, the CG-rich segment forms the same duplex as in the knotted construct, but formation of the AT-rich duplex cannot drive threading because the bulky dsDNA handles prevent the strand from passing through the loop. The molecule therefore folds into a pseudoknot in which the apparent strand crossing is stabilized only by base pairing rather than by a true topological entanglement.

While the folded knot and pseudoknot are very similar in structure, mechanical unfolding will reveal their nanomechanical and topological differences. With all secondary structures obliterated by mechanical pulling, the knot topology is expected to remain within the ssDNA region in the knot construct (Fig.1d), resulting in a shorter contour length compared to the fully unravelled ssDNA from the pseudoknot construct (Fig. 1e).

### Distinct nanomechanical behaviours of ssDNA knot and pseudoknot

We investigated the nanomechanics of our sequence-identical ssDNA knot and pseudoknot by single-molecule force spectroscopy using dual-trap optical tweezers (Fig. 2a). The dsDNA used to lock the ssDNA topology also served as handles in the experiment. After capturing a single tether, the molecule was repeatedly extended and relaxed (Fig. 2b), and thousands of cycles were analyzed (Supplementary Fig. 1). Despite sharing identical sequence and duplex interactions, the knot and pseudoknot exhibited distinct folding, unfolding, and mechanical behaviours.

**Figure 2:**
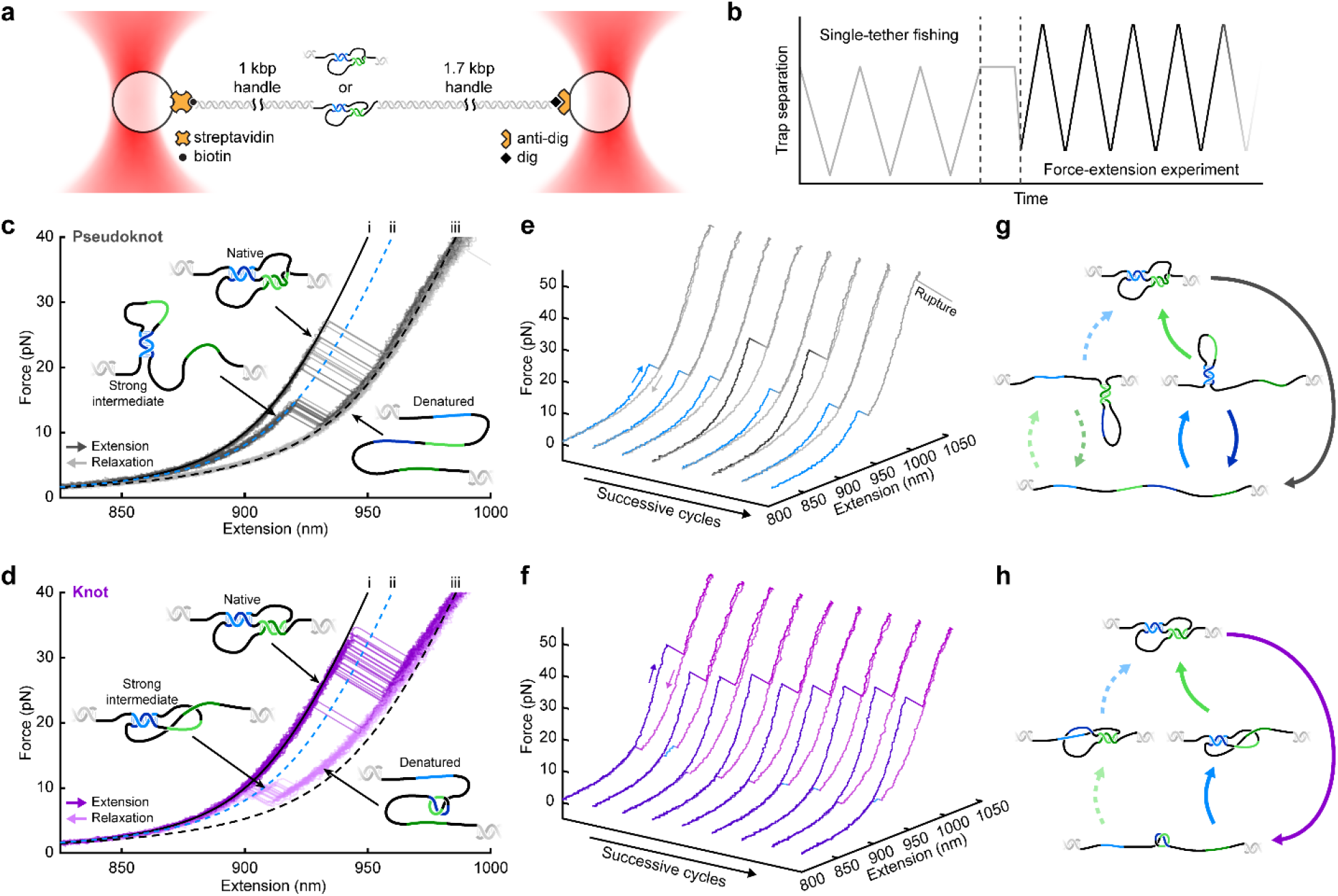
Single-molecule force spectroscopy of ssDNA knot and pseudoknot. **(a)** Schematic of dual-trap optical tweezer experiment. **(b)** After capturing a single tether, the molecule is repeatedly extended and relaxed using a triangular force ramp. **(c)** Representative force-extension curve of a pseudoknot molecule. Dark and light grey traces indicate extension and relaxation, respectively. Dashed curves show extensible polymer models (XWLC for dsDNA and XFJC for ssDNA) corresponding to three states: the native pseudoknot (i), the strong-duplex intermediate (ii), and the fully denatured state (iii). **(d)** Representative force-extension curve of a knot molecule. Dark and light purple traces indicate extension and relaxation, respectively. The same polymer models (i-iii) from (b) are shown for comparison. **(e, f)** Consecutive extension-relaxation cycles for the pseudoknot (e) and knot (f), colour-coded by state. **(g, h)** State transition diagrams of the pseudoknot **(g)** and knot **(h)** based on the force-extension curve measurements. Major folding pathways are indicated by solid arrows and minor pathways by dashed arrows.

Force-extension curves revealed that each tether behaved consistently as either a pseudoknot (Fig. 2c) or a knot (Fig. 2d), with no conversion observed between the two behaviours. This mutual exclusivity indicates that the topological state of the molecule remains locked throughout the experiment. Most molecules (73%) prepared using the knotting process exhibited the knot mechanical behaviour (Supplementary Fig. 2), while the remainder behaved as pseudoknots. Conversely, the majority of molecules (81%) prepared using the pseudoknot process resulted in pseudoknot behaviour, with a small fraction behaving as knot. These minor populations likely arise from imperfect folding yield. In the knotting process, handle ligation may occur before successful strand threading, producing pseudoknot molecules. In the pseudoknot process, incomplete blocking strand binding may allow threading before ligation, producing knotted molecules.

The pseudoknot construct displays two characteristic folding transitions (Fig. 2c). When both duplexes are hybridized (native pseudoknot, i), the CG-rich and AT-rich segments are pulled in a shearing geometry. Because the AT-rich duplex is mechanically weaker, it ruptures first^26–28^ at forces of approximately 20-30 pN, producing an intermediate in which only the CG-rich duplex remains hybridized (strong intermediate, ii). The remaining duplex then rapidly reorients from a shearing to an unzipping geometry and immediately unzips, producing a fully denatured state (iii). During each extension cycle, sufficient force was applied to disrupt both duplexes, ensuring the molecule reached the denatured state. Upon relaxation, however, refolding occurred through two different pathways. Approximately half of the cycles returned to the native pseudoknot at low force, while the remaining cycles refolded only the CG-rich duplex, producing the strong intermediate (ii). In these cases, the subsequent extension began from this intermediate state, resulting in an unfolding transition at a lower force and with a shorter transition distance than unfolding from the native pseudoknot (Fig. 2e).

In contrast, the knot construct exhibits a single unfolding transition at 25-40 pN (Fig. 2d). This transition corresponds to unfolding of the native knot and follows a similar pathway to pseudoknot unfolding, where the weak duplex shears first, followed immediately by unzipping of the strong duplex. The unfolding transition begins from a conformation with a contour length nearly identical to the pseudoknot (i) but unfolds to a state slightly shorter than the denatured pseudoknot (iii). This reduction in contour length arises because knot topology persists even after all base pairs are disrupted, retaining nucleotides within the loops of the knot that shortens the molecule under tension. Upon relaxation, the knot refolds to the native state at 5-10 pN (Fig. 2f,h). Refolding occasionally occurs through a two-step pathway where the strong duplex forms first, producing the strong intermediate, followed by formation of the weak duplex. In contrast to the pseudoknot, these refolding transitions occur at forces readily detectable in the experiment, indicating that the knot topology substantially increases the refolding rate of the molecule.

Although the pseudoknot and knot constructs share the same three stable states – native, strong intermediate, and denatured, the force-dependent transition rates between them differ substantially. In both topologies, refolding involves competition between formation of the strong and weak duplexes. The strong duplex refolds more rapidly because of its greater thermodynamic stability and smaller loop between its complementary segments (Fig. 2g,h). As a result, the dominant folding pathway proceeds through the strong intermediate. Less frequently, the weak duplex folds first (Fig. 2g,h; dashed arrows), producing a weak intermediate state (Supplementary Fig. 3, Supplementary Fig. 4). Regardless of the order of duplex formation, both topologies ultimately fold into the native state. Together, these results show that molecular topology alters both the unfolding pathway and the refolding kinetics of the same DNA sequence. We next quantify how topology modifies the unfolding force, transition distance, and refolding behaviour to define the nanomechanical signatures of nucleic acid knots.

### Mechanical signatures of a ssDNA with knot topology

To identify mechanical signatures of nucleic acid knots, we quantitatively compared the unfolding and refolding behaviour of the sequence-identical pseudoknot and knot constructs (Fig. 3). Because both molecules share the same primary sequence and base-pairing interactions, any differences in their mechanical response arise solely from topology (Supplementary Figs. 5-7). Three measurable features distinguish the knot ssDNA from the pseudoknot: a higher unfolding force, a shorter unfolding distance, and faster refolding kinetics.

**Figure 3:**
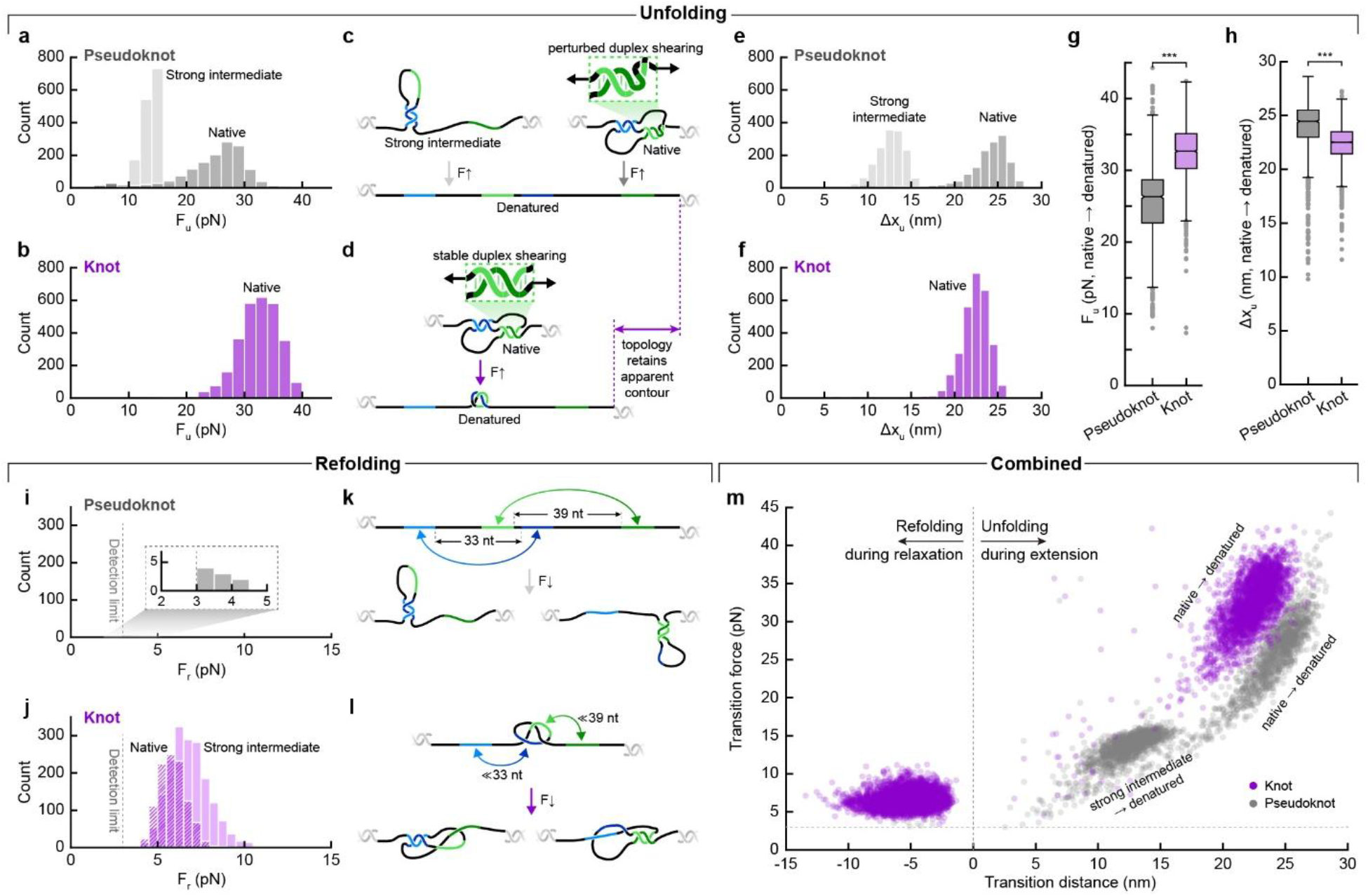
Nanomechanical signatures of a ssDNA knot. **(a, b)** Distributions of unfolding forces (*F*_*u*_) for transitions into the denatured state for the pseudoknot (a) and knot (b) constructs. The pseudoknot exhibits two populations corresponding to unfolding from the native state and from the strong intermediate, whereas the knot unfolds predominantly from the native state. **(c, d)** Schematics illustrating the unfolding mechanisms of the pseudoknot (c) and knot (d). **(e, f)** Distributions of unfolding distances (Δ*x*_*u*_) for the pseudoknot (e) and knot (f). **(g, h)** Comparison of native-to-denatured transitions for the pseudoknot and knot constructs showing higher unfolding force (g) and shorter unfolding distance (h) for the knot. **(i, j)** Distribution of refolding force (*F*_*r*_) for transitions from the denatured state for the pseudoknot (i) and knot (j). Refolding transitions are rarely observed for the pseudoknot above the experimental detection limit (∼3 pN), whereas the knot refolds at higher forces and often proceeds through the strong intermediate. The inset in (i) shows all detected pseudoknot refolding events. **(k, l)** Schematic illustrating the effect of separation between duplex interactions on refolding kinetics. Large loops in the pseudoknot slow duplex formation (k), whereas the knot reduces the available search space and accelerates refolding (l). **(m)** Transition *F*_*u/r*_ vs. Δ*x*_*u/r*_ to and from the denatured state. Knot (purple) and pseudoknot (grey) transitions form distinct clusters of the phase space, providing a mechanical fingerprint of knot topology.

The first signature is the force required to fully unfold the native structure (*F*_*u*_). The unfolding force distribution of the pseudoknot shows two populations corresponding to unfolding from the native pseudoknot and from the strong intermediate (Fig. 3a). In contrast, the knot exhibits a single dominant unfolding transition from the native state (Fig. 3b). When comparing only transitions from the native state to the fully denatured state, the knotted construct unfolds at significantly higher force (33 pN) than the pseudoknot (27 pN, Fig. 3g). This increase arises because both duplex segments remain stably hybridized in the threaded configuration of the knot, whereas in the pseudoknot the unthreaded geometry perturbs the weak duplex under tension (Fig. 3c,d). A knot-stabilized higher unfolding force therefore represents the first mechanical signature of a nucleic acid knot.

A second signature emerges from the extension change associated with unfolding. The unfolding distance Δ*x*_*u*_, defined as the extension difference between native and denatured states at the point of rupture, is larger for the pseudoknot than for the knot (Fig. 3e, f, h). Even though the knot unfolds at a higher force, its Δ*x*_*u*_ is shorter because the topological entanglement persists after base pairs rupture, retaining a looped configuration that shortens the molecule under tension (Fig. 3c, d). Thus, a reduced contour length provides a second mechanical signature of a nucleic acid knot.

The third signature arises during relaxation of the stretched molecule. Refolding transitions from the denatured state are rarely observed for the pseudoknot above the experimental detection limit of ∼3 pN (Fig. 3i), consistent with the slow nucleation kinetics expected for the large loops connecting its duplex segments (Fig. 3k)^27^. In contrast, the knotted construct refolds at substantially higher forces (*F*_*r*_ at 5-10 pN), often through a two-step pathway involving formation of the strong intermediate followed by the native state (Fig. 3j,l). The knot topology effectively reduces the conformational search space for duplex formation, bringing complementary segments into closer proximity and accelerating refolding. The increased refolding force, therefore, constitutes the third nanomechanical signature of a nucleic acid knot.

Plotting all detected transitions into and out of the denatured state as a function of transition forces and distance reveals distinct clusters corresponding to the two topologies (Fig. 3m). Unfolding transitions of the knot occur at higher force and shorter distance than those of the pseudoknot, while rapid refolding transitions are observed only for the knot construct regardless of loading rate (Supplementary Figs. 8-9). Together, these force-distance-kinetics differences define a mechanical signature that distinguishes knotted ssDNA from its pseudoknotted counterpart.

### Force-dependent tightening of a ssDNA knot

To examine the behaviour of the knot in the absence of secondary structure, we analyzed the denatured states of the knot and pseudoknot (Supplementary Fig. 10). Force-extension density maps of the denatured pseudoknot and knot show narrow extension distributions across the applied force range (Fig. 4a,b). The denatured knot consistently exhibits a shorter extension than the pseudoknot (Fig. 4c). Because the two constructs share identical sequence and handles, this difference arises solely from topology. The difference in mean extension therefore defines the contour length retained by the knot (*L*_*knot*_, Fig. 4c). Across the measured force range (>7 pN), the knot remains shorter than the pseudoknot, indicating that the topological entanglement persists after all base pairs are disrupted. Yet *L*_*knot*_ varies continuously with applied force (Fig. 4d).

**Figure 4:**
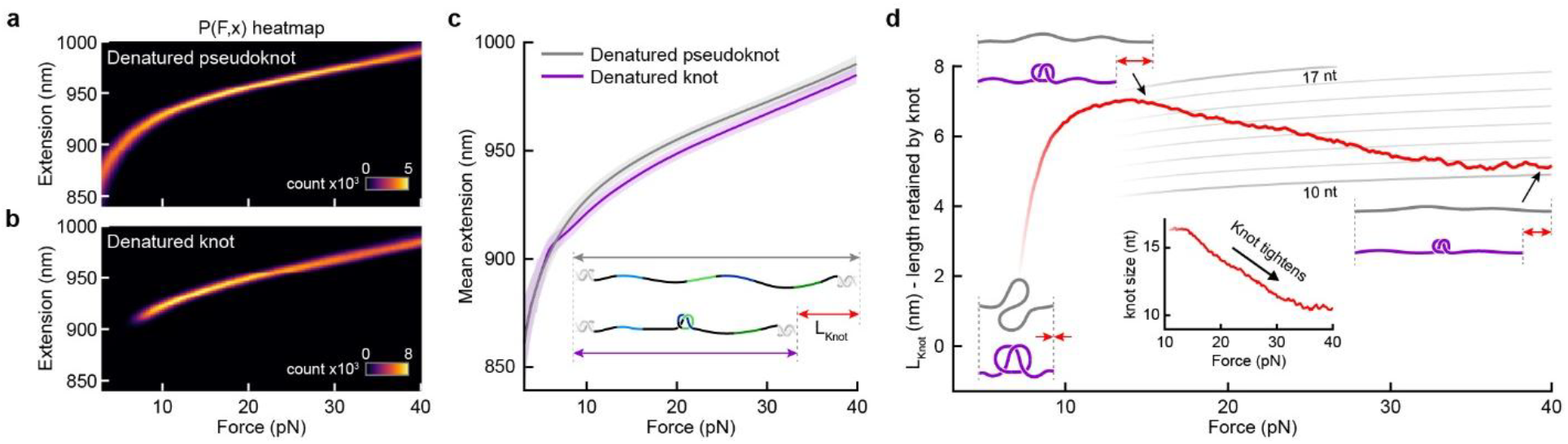
Force-dependent tightening of a ssDNA knot. **(a, b)** Force-extension density maps of the denatured state of the pseudoknot (a) and knot (b) constructs. The denatured knot population deplete below 7 pN due to refolding. **(c)** Mean molecular extension as a function of force for the pseudoknot and knot constructs. Shaded regions indicate the standard deviation. The length retained by the knot (*L*_*knot*_) is defined by the difference in mean extension. **(d)** Length retained by the knot (*L*_*knot*_) as a function of force. The standard error of the mean is less than the line width. Grey lines show the predicted extension of ssDNA segments differing by a single nucleotide based on the XFJC model. Inset: estimated number of nucleotides (nt) within the knot as a function of force.

At low tension, the knot remains loose, and the extension difference between the two constructs is small. In this regime, the force-extension behaviour is dominated by the entropic elasticity of the loose ssDNA, while the knot occupies a relaxed conformation due to low tension (see schematic, Fig. 4d). Consequently, the topological constraint contributes only weakly to the overall extension.

At the high-force regime, the molecule can be modelled as two segments in series: a stretched ssDNA outside the knot and a compact knot. The external strand follows the entropic elasticity of ssDNA, whereas the tight knot resists further tightening because the polymer backbone within the knot must bend sharply. Tightening therefore becomes both entropically and enthalpically unfavourable as the nucleotides composing the knot lose configurational freedom while simultaneously increasing in bending energy. As force increases, nucleotides (nt) are progressively pulled from the knot into the stretched ssDNA segment outside, effectively shrinking the knot size and reducing *L*_*knot*_. By comparing the measured *L*_*knot*_ with the predicted contours of ssDNA (Fig. 4d), we estimate the number of nucleotides contained within the knot as a function of force (Fig. 4d inset). From the transition near ∼13 pN to the experiment limit of 40 pN, the tight knot contracts from ∼17 to ∼10 nt.

## Methods

### Pseudoknot and knot folding processes

Full custom DNA oligo sequences are available in Supplementary Table 1 (Integrated DNA Technologies, Coralville, IA, USA). The two long handles were built separately. The left handle was made by polymerase chain reaction (PCR) amplification using Taq DNA polymerase (FroggaBio, Toronto, Canada) of a 1 kbp region of the pBR322 plasmid (New England Biolabs, Whitby, Canada) incorporating a 5’ biotin modification on the forward primer. This construct was subsequently digested with PspGI (New England Biolabs, Whitby, Canada) at 75 °C for 1 hour to create sticky ends for subsequent ligation to the knot-forming sequence. The 1.7 kbp right handle was PCR amplified from the lambda genome (New England Biolabs, Whitby, Canada) while also incorporating a 5’ modified digoxigenin. A sticky end was created through digestion with TspRI at 65 °C for 1 hour. Both constructs were purified using a Monarch PCR & DNA Cleanup Kit (New England Biolabs, Whitby, Canada).

In the knot-forming process, the knot-forming sequence was added at 0.4 µM to a buffer of 50 mM NaCl. The sample was heated in a thermocycler to 95 °C for 5 minutes, then cooled immediately to 70 °C. This was followed by slow cooling at 1 °C/min until 20 °C was reached. The extra magnesium condition (Supplementary Fig. 1) was performed identically, except with an additional 5 mM of MgCl_2_ in the sample. The pseudoknot-forming process was treated identically and prepared simultaneously, except that 1.6 µM blocking strand was added prior to the initial heat denaturation.

Ligation was performed overnight at room temperature with relative molar ratios 1:1:0.8 of the left handle, right handle, and knot/pseudoknot construct, respectively, using T4 DNA ligase (New England Biolabs, Whitby, Canada). This was then run on a 0.8% agarose gel pre-stained with gel-green (Biotium, Fremont, CA, USA), and the resulting full-length construct was gel-purified using a Monarch DNA Gel Extraction Kit (New England Biolabs, Whitby, Canada). In the pseudoknot-forming process, the purified sample was diluted 10X in water, and the blocking strand was dislodged by heating to 62°C for 5 minutes, followed immediately by an additional DNA cleanup to remove it. Following cleanup, the resulting stable pseudoknot and knot were stored at -20°C.

### Optical tweezers experiments

The stable pseudoknot and knot samples were removed from the freezer for optical tweezers experiments. Protein-A-coated 1050 nm diameter polystyrene beads (Bangs Labs, Fishers, IN, USA) were incubated in 10 µg/mL anti-digoxigenin (Roche Diagnostics, Mannheim, Germany) to create anti-digoxigenin beads. Excess anti-digoxigenin was then washed out. Approximately 0.2 ng/µL of the stable construct was then incubated with streptavidin-coated 1050 nm diameter polystyrene beads (Bangs Labs, Fishers, IN, USA) for 30 minutes. Both the anti-digoxigenin and streptavidin beads were then washed with a buffer of 50 mM NaCl, 10 mM Tris-HCl (pH 8.0) and 5 mM MgCl_2_. This same buffer was used for the subsequent optical tweezers experiments, with the addition of an oxygen scavenger system containing 20 mM protocatechuic acid (Sigma-Aldrich, Oakville, Canada) and 50 nM protocatechuate-3,4-dioxygenase. For the optical tweezers experiments, three separate inputs into a custom flow chip were prepared: the streptavidin-DNA beads as described above, the anti-digoxigenin-coated beads, and a bead-free tube of the trapping buffer. The custom flow chip was made of two no. 1 coverslips (Thermo Fisher Scientific, Nepean, Canada) sandwiching CNC-cut parafilm to create channels shaped like a trident. The coverslip and parafilm sandwich was melted and, after cooling, formed a sealed chamber with three syringe pump-controlled laminar inputs and a single output. Syringe pumps were used to refresh the trapping buffer and to introduce new beads between individual bead-pair experiments, but no flow was used during data acquisition.

A streptavidin-DNA bead was held in one trap of the dual-trap optical tweezers, while the other held an anti-digoxigenin bead. All single-molecule experiments were performed at approximately 26°C on a custom-built dual-trap optical tweezer as described in our previous work^29^. Individual bead calibration data were acquired at 100 kHz, with subsequent dynamic force spectroscopy data recorded at 10 kHz. The beads were steered close together until a tether was formed. Data collection was performed in repeated cycles of extension and relaxation, steered by a triangle wave at either 252.5 nm/s (slow) or 1010 nm/s (fast), until the tether ruptured. The traps were always steered the same total distance at each rate. The low-end force was kept below 0.5 pN, while the high-end was above 40 pN but below the DNA overstretching regime.

### Single Molecule Data Analysis

Data processing was performed using custom Matlab code available alongside complete datasets on Zenodo^30^. In brief, each bead was individually calibrated using well-established theory^31^, the force measured by both traps were averaged to minimize the impacts of optical instability. In cases of multiple tether formation, poor-quality bead calibration, fluidic instability, or debris entering the trap, the data were removed. Tethers were manually classified by their behaviour into either pseudoknot or knot-like behaviour. Unfolding and refolding transitions were detected using the *findchangepts* function, with the initial and final states of each transition determined by comparison with the modelled XWLC states for native, weak hairpin, strong hairpin, and denatured (Supplementary Tables 2,3). Some manual reclassifications were performed to correct misclassified states and remove any detected “transitions” that were just experimental noise. Missing low-force transitions were inferred from the observed states immediately before and after the unobservable low-force region. Detection of short-lived states lasting less than 1 ms was not performed, as we required at least 10 data points per state to assign it with a reasonable degree of certainty. Heat maps for the most likely extension at a given force were generated by first classifying each point in the force extension curve with a state and a pulling direction. States were found by setting everything before a transition to its initial state and everything after a transition to its final state. Inferred transitions (and thus inferred states) were not used for this analysis. A buffer region of 10 points before and 10 points after a detected transition was not classified in any state and thus omitted from the heat map. Only data in the denatured state and in the relaxation steering direction were used for Fig. 4. The number of nucleotides in the knot was estimated by subtracting the mean extension at each force for the knotted construct from the mean extension at each force for the pseudoknot. For each force, this difference was interpolated against the extension of ssDNA of sequence lengths from 1-99 nt at that force. The resulting estimated number of nucleotides was reported.

## Data Availability

The complete single-molecule data and analysis code are available in the Zenodo repository^30^.

## Notes

### Competing Interest Statement

The authors have declared no competing interest.

### Funding

- Natural Sciences and Engineering Research Council: RGPIN-2024-04352
- Canada Research Chairs: CRC-2020-00143
- Michael Smith Health Research BC: SCH-2020-0559
- Canada Foundation for Innovation: 35492, 40086, 41104

