## Supplementary Info for "Mechanical signatures of nucleic acid knot topology"

### Supplementary Data Figures & Tables

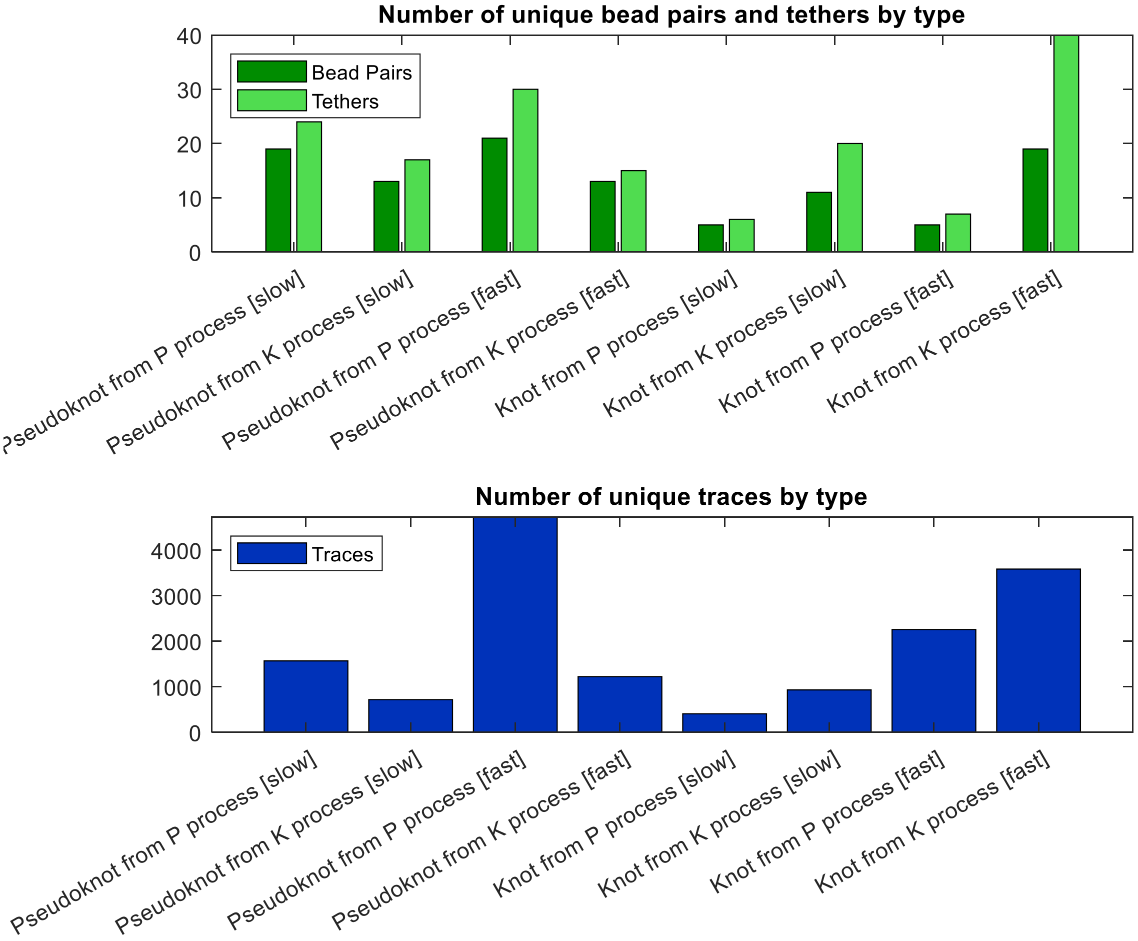


**Supplementary Figure 1.** The number of individual bead pairs and tethers observed from each folding process (K or P meaning knot forming process or pseudoknot forming process, see Fig. 1), classified into pseudoknot or knot behaviour at each pulling rate (top). Some beads were re-fished, resulting in multiple tethers from the same pair of beads which could plausibly be the same single molecule tether. The number of individual traces (extension and relaxation are counted as one trace each) categorized same as above (bottom).

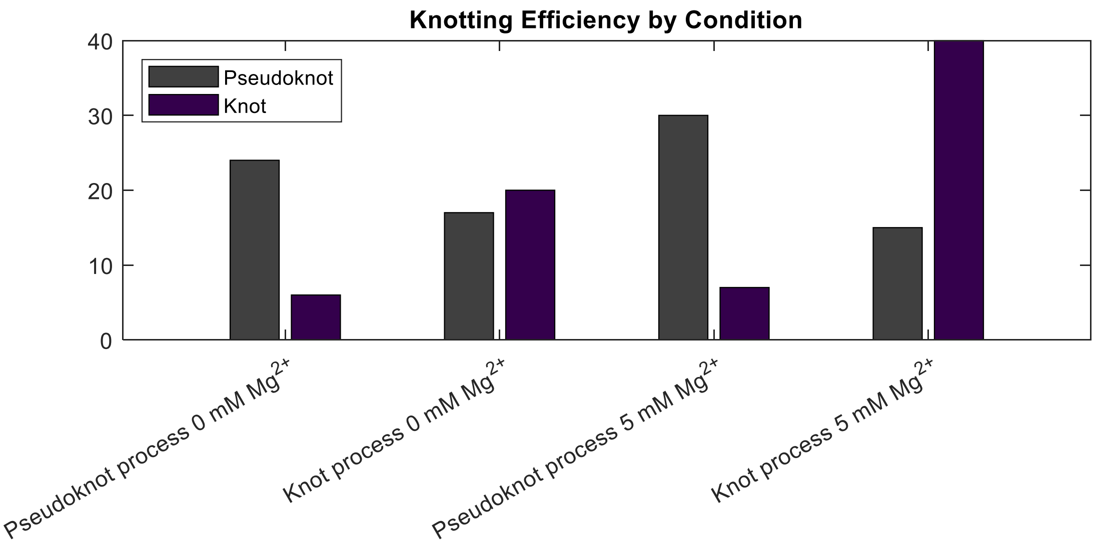


**Supplementary Figure 2.** Efficiency of pseudoknot and knot processes at two different salt concentrations (data categorized by single-molecule behaviour). The knotting efficiency is increased with a higher $Mg^{2+}$ concentration, while having minimal impact on the pseudoknot process success rate.

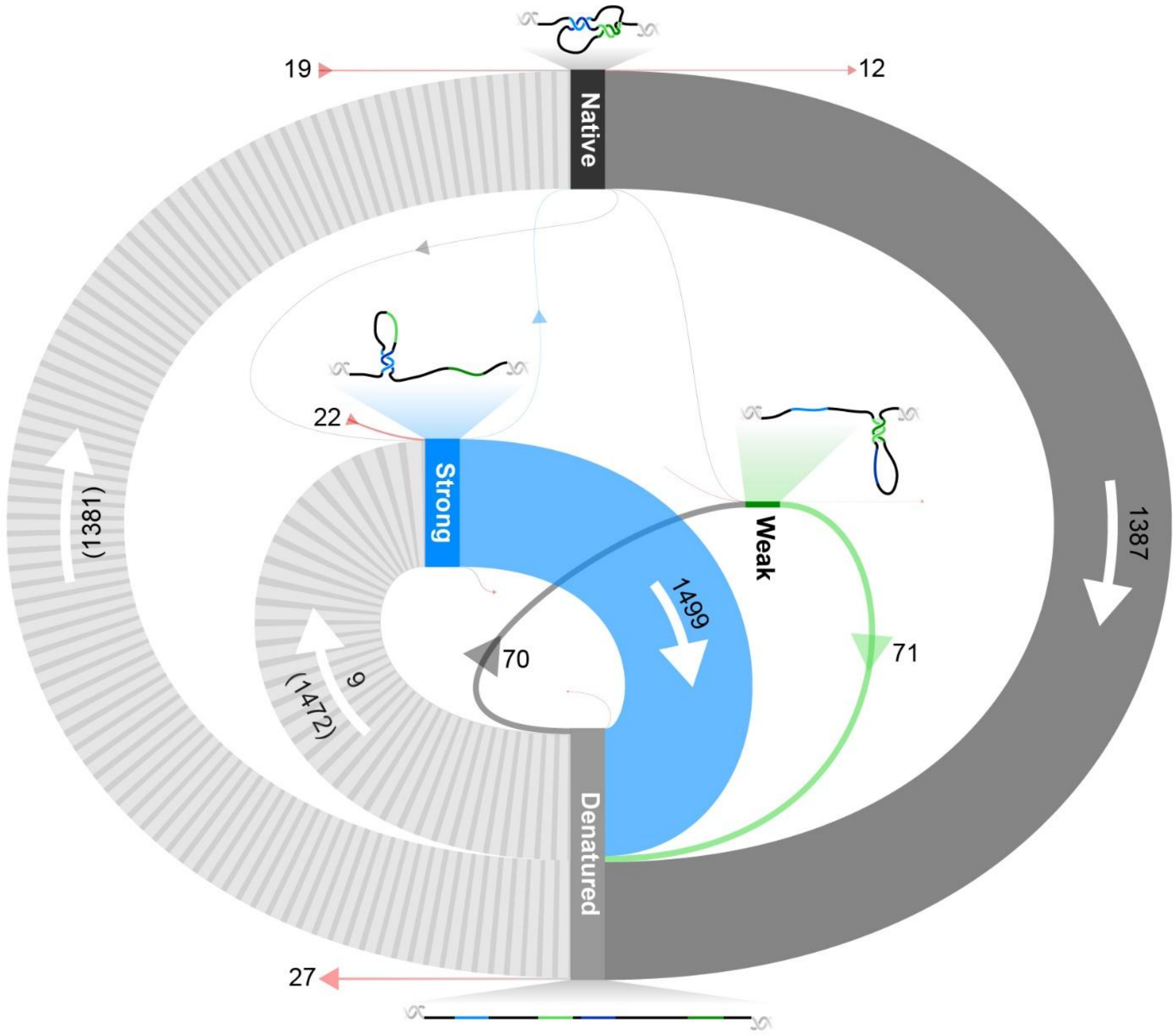


**Supplementary Figure 3:** Each unfolding/refolding transition has an initial state and final state. Transitions from all molecules at the fast steer rate in a state flow map for the pseudoknot are presented here. Arrows indicate the direction of the transition, line width quantitatively represents the number of transitions, and flows are coloured by the initial state of the transition. Inferred transitions between states where the refolding event occurred at such low force to not be directly observable are shown here with stripes. Number of transitions are written on each line (inferred transitions are in brackets, observed transitions are not in brackets). Refolding events from the denatured state to the native state are not thought to occur directly via this pathway; however, because they occur at forces below our observable range we cannot quantitatively determine which intermediate they first folded into. It is known that they subsequently base paired the other interaction to achieve the native state. Regardless of the refolding path, the same native state is achieved. The first observed state of each tether is shown as a red arrow from empty space into a state, while final rupture of a tether is shown as a red arrow out of the state into empty space. State flows with less than 10 transitions (observed and inferred combined) have text labels omitted for clarity.

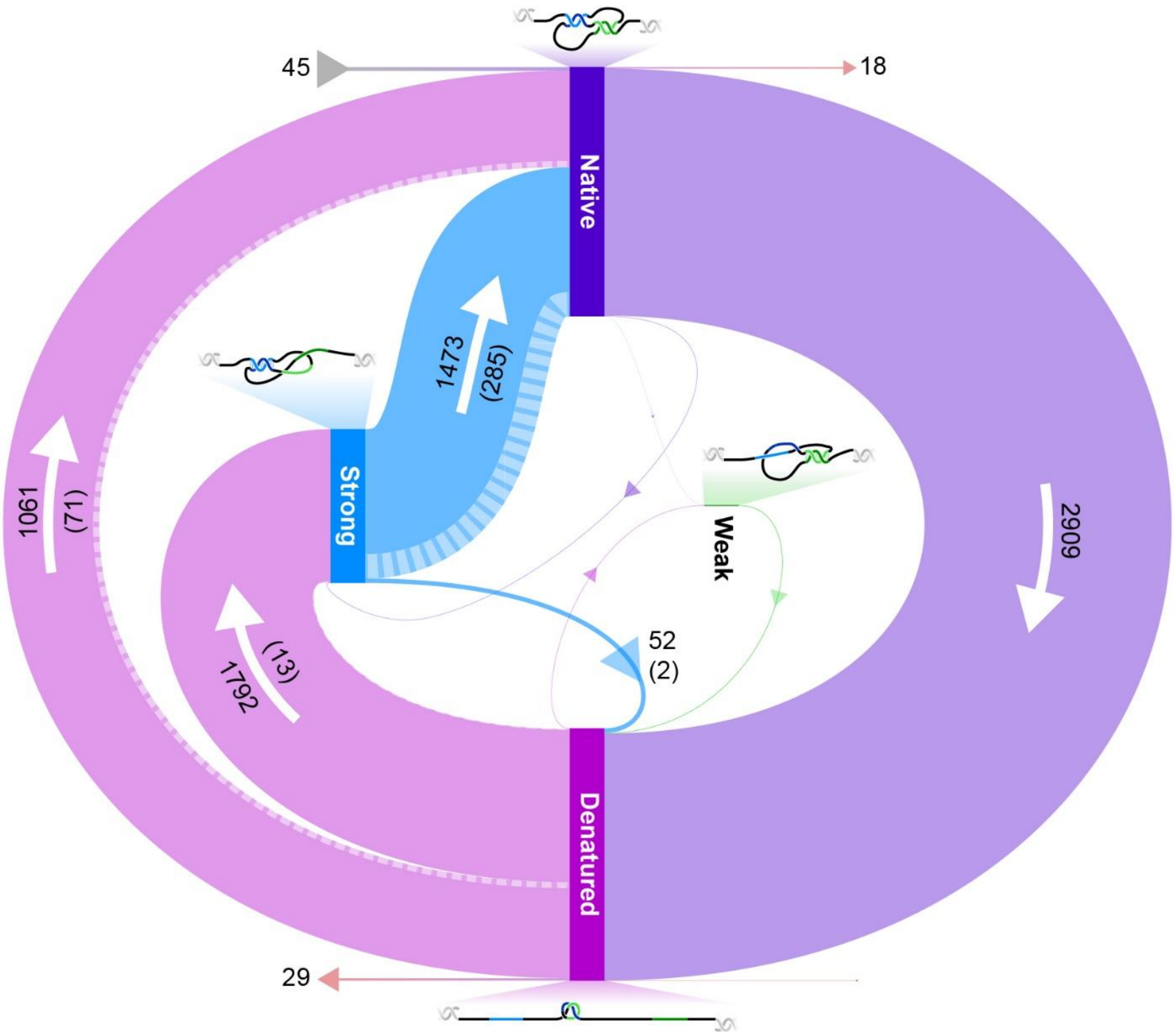


**Supplementary Figure 4:** Transitions from all molecules at the fast steer rate in a state flow map for the knot are presented here. Each unfolding/refolding transition has an initial state and final state. Arrows indicate the direction of the transition, line width quantitatively represents the number of transitions, and flows are coloured by the initial state of the transition. Inferred transitions between states where the refolding event occurred at such low force to not be directly observable are shown here with stripes. Number of transitions are written on each line (inferred transitions are in brackets, observed transitions are not in brackets). Refolding events from the denatured state to the native state are not thought to occur directly via this pathway but are hypothesized to occur via either the strong or weak intermediate with such short-lived lifetimes that we cannot confidently determine which intermediate pathway they first folded into on their towards the native state. The first observed state of each molecule is shown as a red arrow from empty space into a state, while final rupture of a tether is shown as a red arrow out of the state into empty space. State flows with less than 10 transitions (observed and inferred combined) have text labels omitted for clarity.

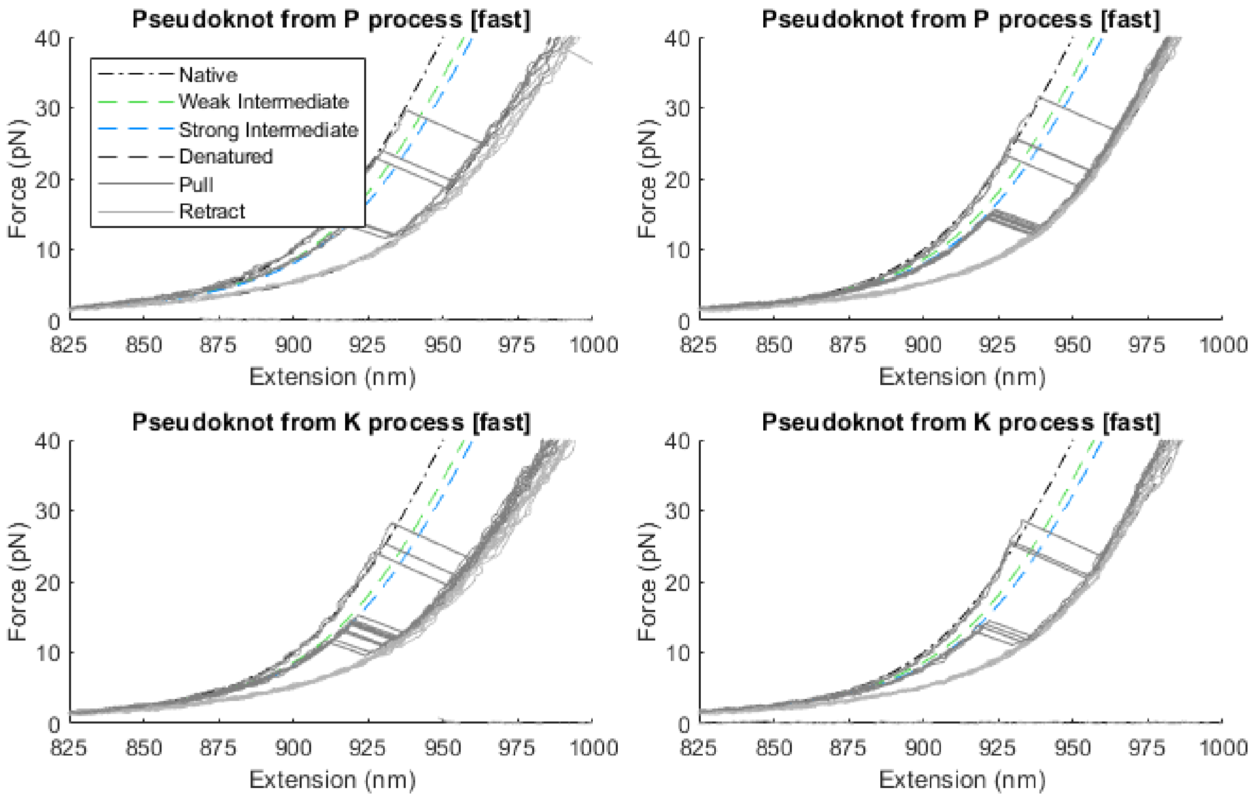


**Supplementary Figure 5.** Two representative FEC examples of pseudoknots steered at the fast steering rate from both the pseudoknot (P, top) and knot (K, bottom) processes.

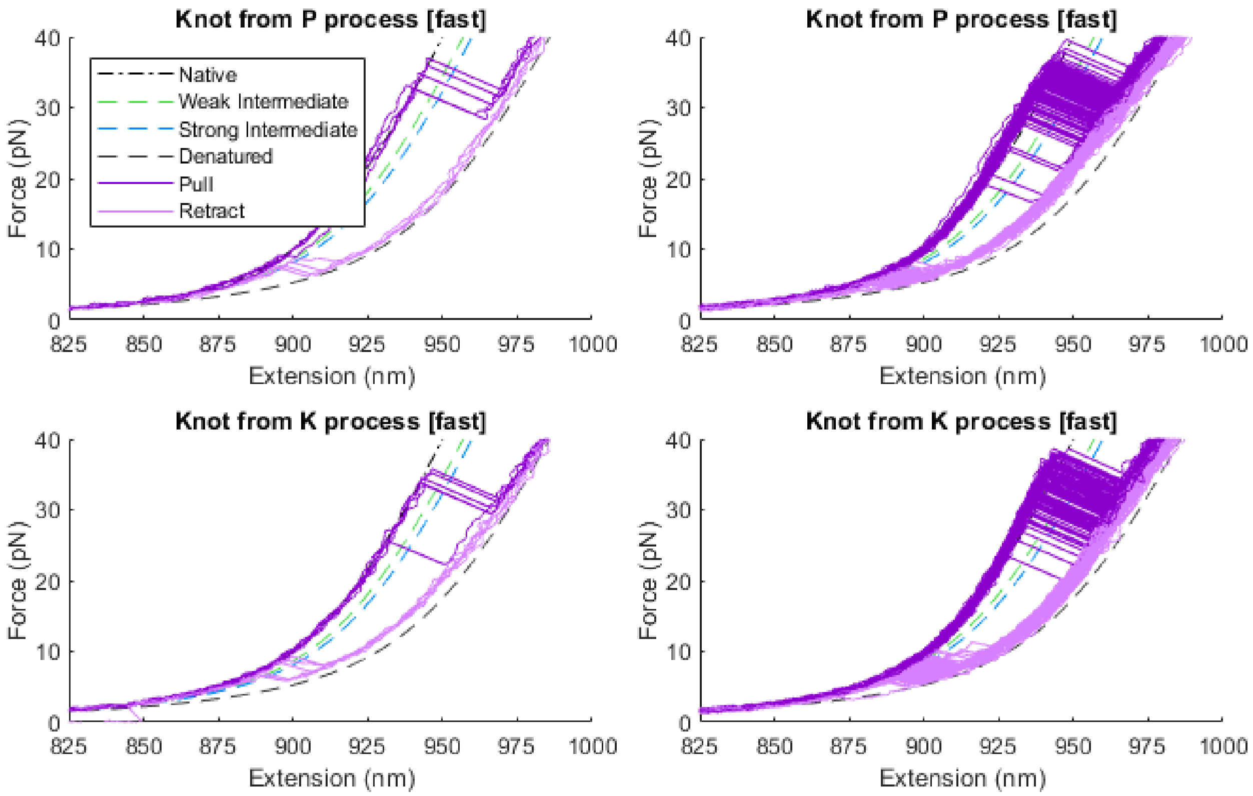


**Supplementary Figure 6.** Two representative FEC examples of knots steered at the fast steering rate from both the pseudoknot (P) and knot (K) processes.

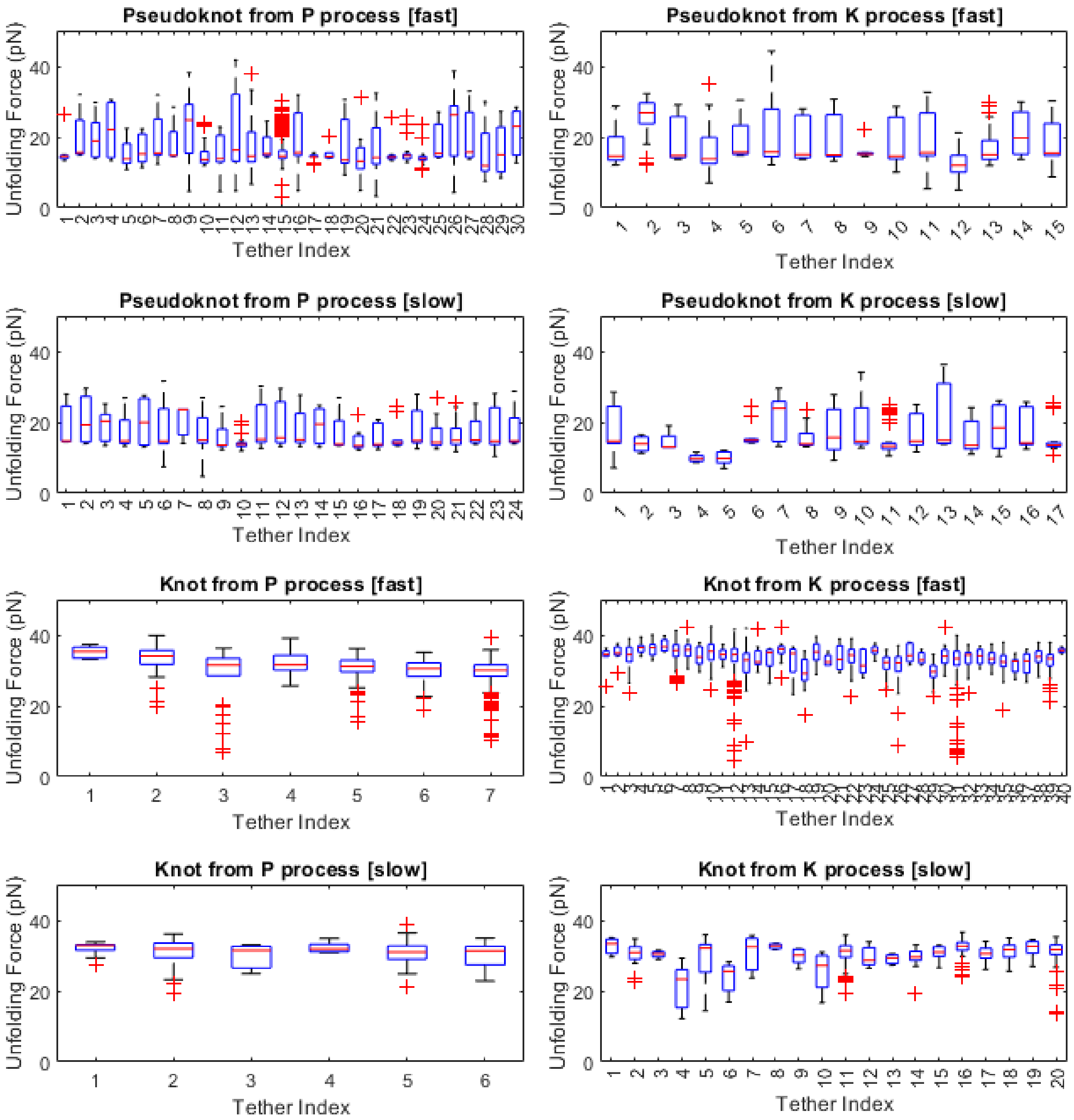


**Supplementary Figure 7:** Observed unfolding forces sorted by individual tether. Pseudoknot formation process (P process, right) and knot formation process (K Process, right) are separated to compare behaviour from each method. Most variation occurs between tethers rather than between processes (a knot from the K process behaves the same as a knot from the P process). Tether-tether variance are hypothesized to originate from the underlying distribution of varying bead size used in the optical tweezers experiments.

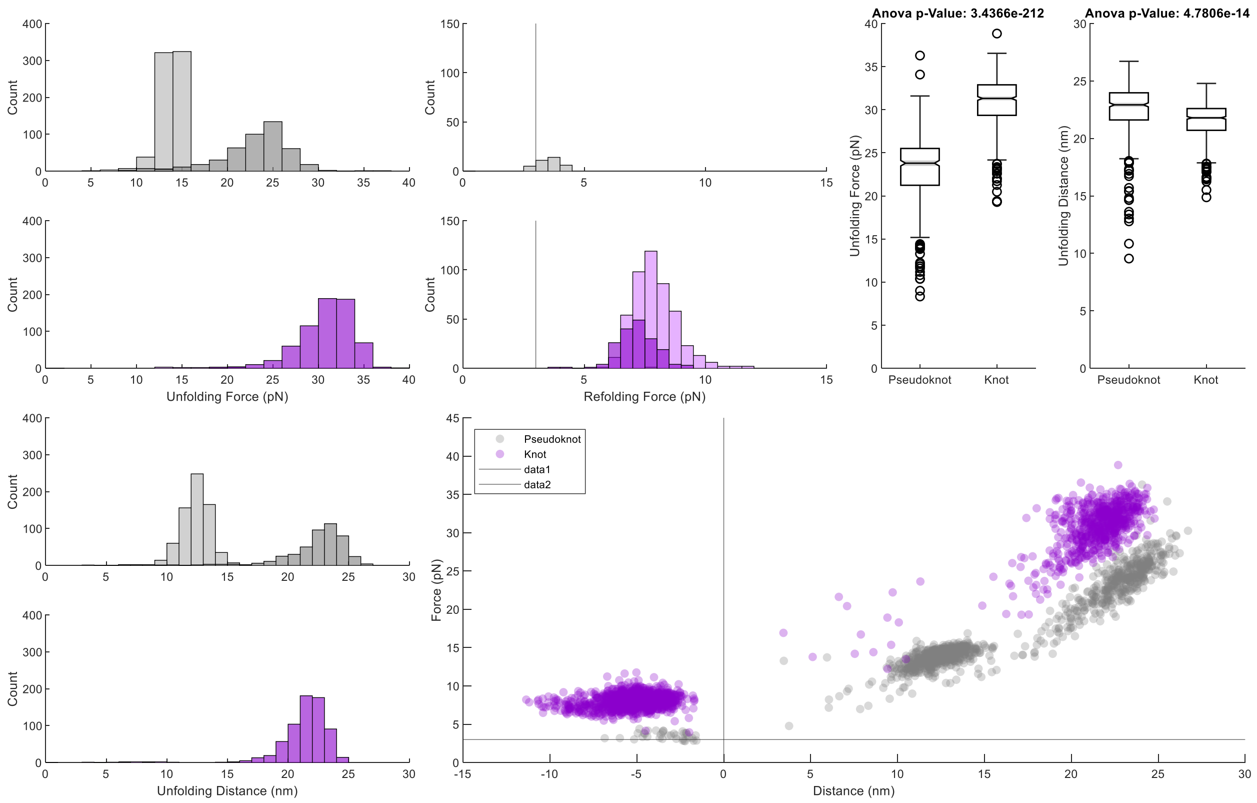


**Supplementary Figure 8.** The same plotted information as for Figure 3, but the slow steering rate data. Changes in transitions occur as expected, with slightly more refolding events observed in the pseudoknot due to the slower steering alongside predictable changes to the unfolding transitions expected with a slower steering rate.

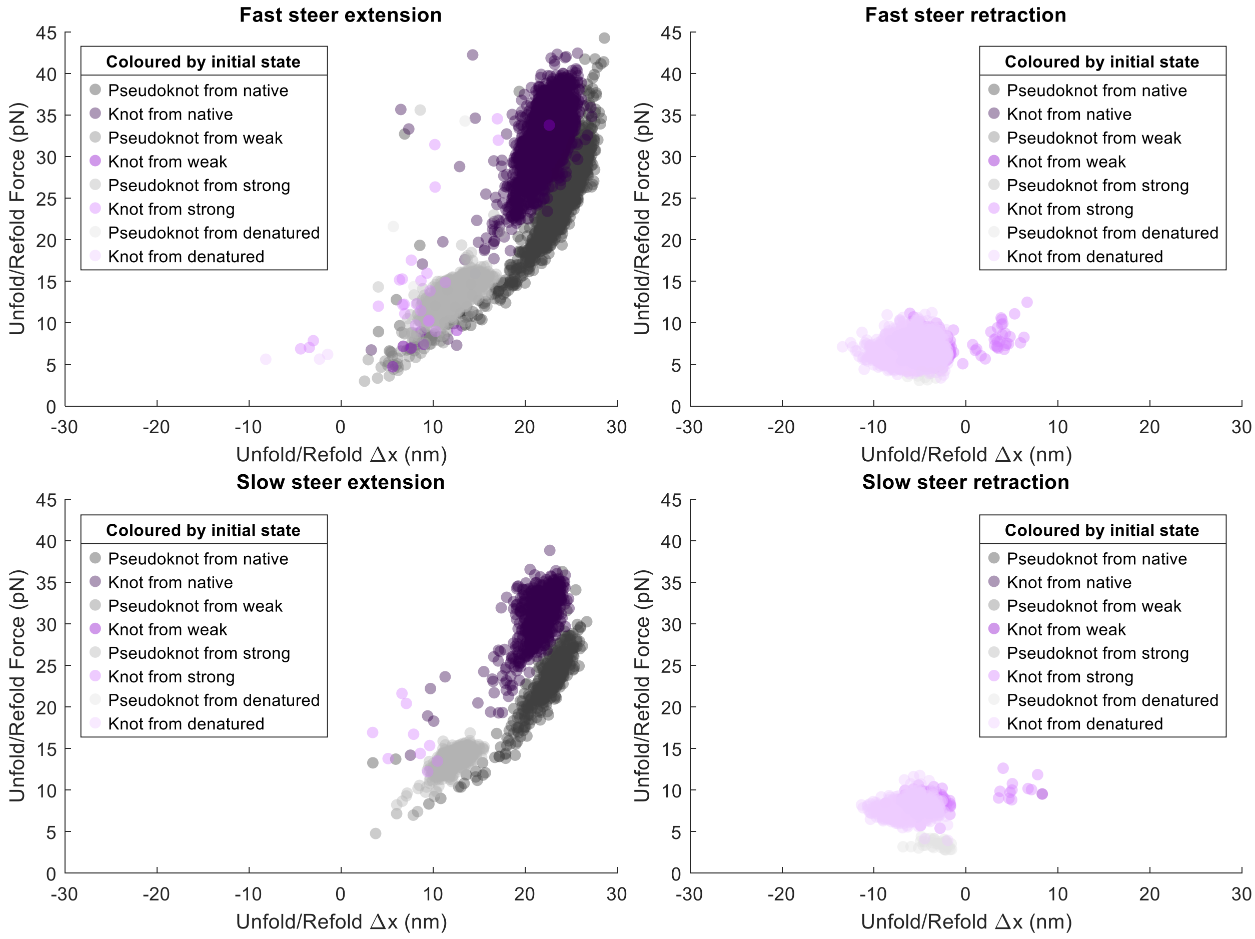


**Supplementary Figure 9.** Transition force and Δx scatterplots of all detected unfolding and refolding events, coloured by their initial state and grouped into different plots by the steering rate (fast, top or slow, bottom) and direction.

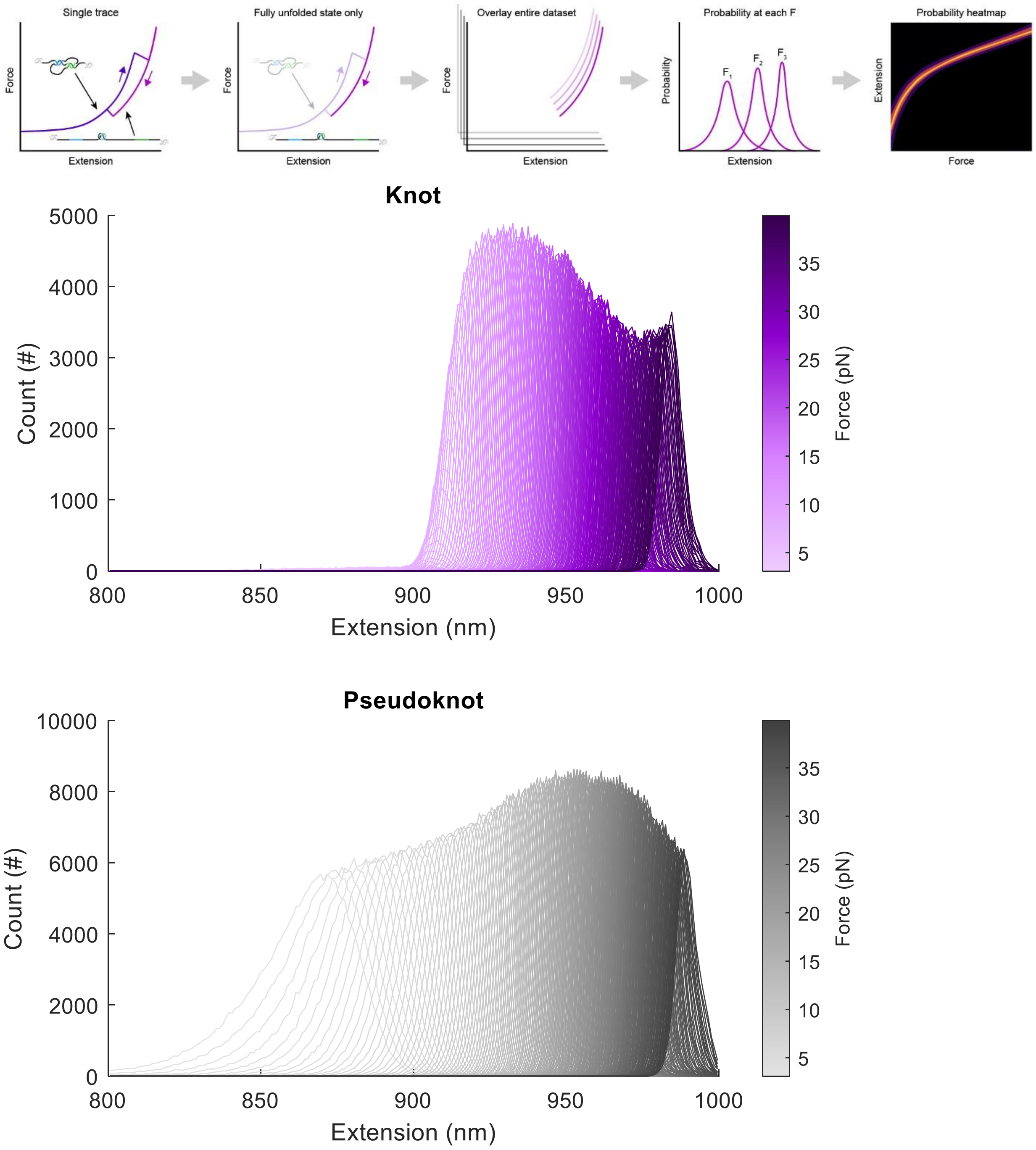


**Supplementary Figure 10:** (Top) Schematic of data processing used to create main text Figure 5. (Middle and Bottom) Individual extension distributions of the denatured state for each force, used to create force-dependent density maps of both topologies. The sample sizes are not even at each force because of transitions into and out of the denatured state occurring at different rates depending on the topology.

**Supplementary Table 1:** Data table of the custom ssDNA oligos used in this study. All oligos were ordered from Integrated DNA Technologies. The following codes, matching the order codes of Integrated DNA Technologies were used in the order: /5BiosG/ for 5' biotin modification; /5DigN/ for 5' digoxigenin modification; and /5Phos/ for 5' phosphate modification for the purposes of subsequent ligation. Nucleotides in blue from the strong interaction, while those in green form the weak interaction.

| Name | Sequence (5' to 3') | Purification |
|---|---|---|
| 1 kbp handle forward primer | /5BiosG/ CAGAAGTGGTCCTGCAACT | HPLC |
| 1 kbp handle reverse primer | CAAGCCTATGCCTACAGCAT | Standard desalting |
| 1.7 kbp handle forward primer | /5DigN/ GGGCAAACCAAGACAGCTAA | HPLC |
| 1.7 kbp handle reverse primer | CGTTTTCCCGAAAAGCCAGAA | Standard desalting |
| Knot forming sequence | /5Phos/CCTGG TTT CGGCGTCCTGCG TTTTTTTTTTTTTTTTTT GTAATGATTACG TTT CGCAGGACGCCG TTTTTTTTTTTTTTTTTTTTTTTT CGTAATCATTAC TTT CCCACTGGC | Ultramer (Standard) |
| Blocking sequence | CGGCGTCCTGCG AAA CGTAATCATTAC | Standard desalting |

**Supplementary Table 2:** Parameters and models used for polymer modelling.

| Polymer & Model | Parameter | Value |
|---|---|---|
| ssDNA<br>Extensible freely jointed chain | Kuhn length | 2.2 nm |
| | Persistence length | 1.1 nm |
| | Stretch modulus | 800 pN |
| | Contour Length | 0.49 nm/nt |
| dsDNA<br>Extensible worm like chain | Persistence Length | 50 nm |
| | Stretch Modulus | 1000 pN |
| | Contour Length | 0.34 nm/nt |

**Supplementary Table 3:** Modelled force extension curve states including their various components.

| Name | dsDNA (bp) | ssDNA (nt) | Helix Widths (#) |
|---|---|---|---|
| Native | 2734 | 9 | 0 |
| Weak intermediate | 2710 | 36 | 1 (2 nm) |
| Strong intermediate | 2710 | 42 | 1 (2 nm) |
| Denatured | 2710 | 99 | 0 |
